# Active microbial communities facilitate carbon turnover in brine pools found in the deep Southeastern Mediterranean Sea

**DOI:** 10.1101/2023.11.26.568696

**Authors:** Maxim Rubin-Blum, Yizhaq Makovsky, Eyal Rahav, Natalia Belkin, Gilad Antler, Guy Sisma-Ventura, Barak Herut

## Abstract

Discharge of gas-rich brines fuels productive chemosynthetic ecosystems in the deep sea. In these salty, methanic and sulfidic brines, microbial communities adapt to specific niches along the physicochemical gradients. However, the molecular mechanisms that underpin these adaptations are not fully known. Using metagenomics, we investigated the dense (∼10^6^ cell ml-^1^) microbial communities that occupy small deep-sea brine pools found in the Southeastern Mediterranean Sea (1150 m water depth, ∼22°C, ∼60 PSU salinity, sulfide, methane, ammonia reaching millimolar levels, and oxygen usually depleted), reaching high productivity rates of 685 µg C L^-1^ d^-1^ ex-situ. We curated 266 metagenome-assembled genomes of bacteria and archaea from the several pools and adjacent sediment-water interface, highlighting the dominance of a single *Sulfurimonas*, which likely fuels its autotrophy using sulfide oxidation or inorganic sulfur disproportionation. This lineage may be dominant in its niche due to genome streamlining, limiting its metabolic repertoire, particularly by using a single variant of sulfide: quinone oxidoreductase. These primary producers co-exist with ANME-2c archaea that catalyze the anaerobic oxidation of methane. Other lineages can degrade the necromass aerobically (*Halomonas* and *Alcanivorax*), or anaerobically through fermentation of macromolecules (e.g., Caldatribacteriota, Bipolaricaulia, Chloroflexota, etc). These low-abundance organisms likely support the autotrophs, providing energy-rich H_2_, and vital organics such as vitamin B12.

## 1. Introduction

Brine pools are extreme hypersaline habitats, where oxygen is depleted, methane fluxes are high and sulfur turnover is substantial (Joye et al., 2009). They are found in various seas, including the Gulf of Mexico, the Mediterranean Sea, the Black Sea and the Red Sea (Merlino et al., 2018; Purkis et al., 2022). Along the continental margins, brine pools are formed due to pervasive hydrocarbon seepage and mud volcanism (Joye, 2020). In the abyssal seabed, brines are associated with the deep-sea hypersaline anoxic basins (DHABs), formed by the dissolution of anciently buried evaporites (Cita, 2006; Duarte et al., 2020; Purkis et al., 2022; Wallmann et al., 1997). Mud volcanism is particularly prominent in the eastern Mediterranean Sea (EMS), fueling vast chemosynthetic ecosystems (Bayon et al., 2009; Charlou et al., 2003; Huguen et al., 2009, 2005; Olu-Le Roy et al., 2004; Omoregie et al., 2008; Teske and Carvalho, 2020). DHABs are also widespread in the Mediterranean Sea, as brines accumulate at the seabed of collapsed basins (Cita, 2006; La Cono et al., 2019). In the EMS, the chemical composition of the brines depends on the succession of Messinian evaporite precipitation, as well as dissolution and advection mechanisms (Cita, 2006; De Lange et al., 1990; Vengosh et al., 1998). The eastmost EMS brine pools fed by the Messinian evaporites were recently discovered at the Palmahim Disturbance offshore Israel (Herut et al., 2022).

Diverse bacteria and archaea thrive at brine pools and lakes, driving dark productivity and nutrient turnover (Borin et al., 2009; Daffonchio et al., 2006; Joye et al., 2009; Kormas et al., 2015; Merlino et al., 2018; Nigro et al., 2020; Pachiadaki and Edgcomb, 2022; Pachiadaki et al., 2014; Steinle et al., 2018; Yakimov et al., 2013, 2015; Zhang et al., 2016). These microbes are often adapted to high salinities, as suggested by the high expression of genes involved in the synthesis and transport of organic and inorganic osmoprotectants (Pachiadaki et al., 2014). Brine microbes may form dense (circa 10^6^ cells ml^-1^) and highly productive populations at the brine interface (also known as pycnocline, halocline, or chemocline), whereas in the deeper parts of brines their activity may be limited due to energetic constraints, and lean on fermentation, acetogenesis and methanogenesis (Lazar, 2020; Mwirichia et al., 2016; Pachiadaki and Edgcomb, 2022; Schuchmann and Müller, 2014; Steinle et al., 2018; Yakimov et al., 2013, 2007).

Sulfur redox dynamics is likely the key driver of microbial productivity at the brine interface (Borin et al., 2009). The reduced sulfur compounds originate in the lower brine interface, where sulfate-reducing Desulfobacterota (i.e., Deltaproteobacteria) catalyze sulfate reduction. Rates ranging between 20.6 and 460 µmol kg^−1^ d^−1^ were measured in Urania and Kryos lakes (Borin et al., 2009; Steinle et al., 2018), whereas extremely high sulfate reduction rates of circa 20 mmol kg^−1^ d^−1^ were measured in Bannock basin brines (Daffonchio et al., 2006). Hydrogen turnover may also play a role in fueling the productivity of brine organisms (Joye et al., 2009). Stable isotope analyses of biomarkers and genetic markers indicated that Campylobacterota thiotrophs (i.e., Epsilonproteobacteria) fixed carbon in Kryos hypersaline anoxic lake brines using mainly the reverse tricarboxylic acid (rTCA) cycle (Steinle et al., 2018). Metagenomics suggests that the rTCA cycle was the key mechanism for the fixation of inorganic carbon at the interface of the Thetis Lake (Ferrer et al., 2012). In turn, macromolecules from the primary producers can be recycled by diverse organisms (Zhang et al., 2016).

Methane is another prominent source of carbon and energy in brines, reaching millimolar levels and being produced intrinsically at rates of 169 µmol L^−1^ d^−1^ (Borin et al., 2009). Aerobic and anaerobic removal of methane by type-I methanotrophs Methylococcales and anaerobic methane-oxidizing (ANME) archaea can be substantial at the brine interface (La Cono et al., 2019; Lazar, 2020; Pachiadaki et al., 2014; Steinle et al., 2018). Whereas the ANME-1 clade appears to be dominant at the halocline of Mediterranean DHABs, ANMEs are absent in deeper, highly chaotropic sections of the MgCl_2_-rich brines in athalassohaline DHABs such as Kryos and Hephaestus (La Cono et al., 2019; Steinle et al., 2018).

Here we focus on microbial diversity and functions in thalassohaline brines in the eastmost Mediterranean brine pools found at the Palmahim Disturbance offshore Israel (Herut et al., 2022). These small and shallow pools (one to several meters in diameter, depths less than a few meters) are located at a water depth of circa 1150 m. They are anoxic, methanic and warm (21.6 °C), with maximal salinities of 63.9 PSU (Herut et al., 2022). We quantified prokaryotes and their productivity rates at the brine-water interface and studied the genomic potential of the intrinsic microbes to fix carbon and catalyze nutrient turnover. We aimed to identify metabolic handoffs and biotic interactions in communities from several pools, as well as those overlaying the brine-infused sediments.

## 2. Materials and Methods

### 2.1 Sample collection

the studied brine pools are located at the toe of Palmahim Disturbance, a large scale (∼50 km by 15 km) rotational slide offshore Israel, at a water depth of circa 1150 m (32.22° N 34.18° E). We sampled brines and top core water during two seafloor surveying legs (in April and November 2021), using the University of Haifa Jonah SAAB Seaeye Leopard Remotely Operated Vehicle (ROV) operated onboard Israel Oceanographic and Limnological Research (IOLR) R/V Bat-Galim. Brines were sampled using a Niskin bottle mounted on the ROV and water was collected from sediment cores (**Figure 1**, **Supplementary Video 1**). We collected six samples of brine interface (BP1-5), as well as overlying water from two push cores (PC1 and PC2). BP2 and BP4-5 were collected from a relatively deep 5-m-wide pool, whereas samples BP1 and BP3 were collected from other less deep pools, and contained contaminating sediments from resuspended from the pool’s bottom. Given that the depth of the pools ranged between several decimeters to the order of 1 meter, the putative disturbance of water with the ROV thrusters, and the substantial time needed to fully exchange water in the Niskin Bottle, our samples contain some contamination from overlying water and sediments from the pool bottom. CTD was deployed in the sampled brines, to in-situ measure pressure, salinity and dissolved oxygen (Herut et al., 2022) (**Supplementary Figure S1**).

**Figure 1:**
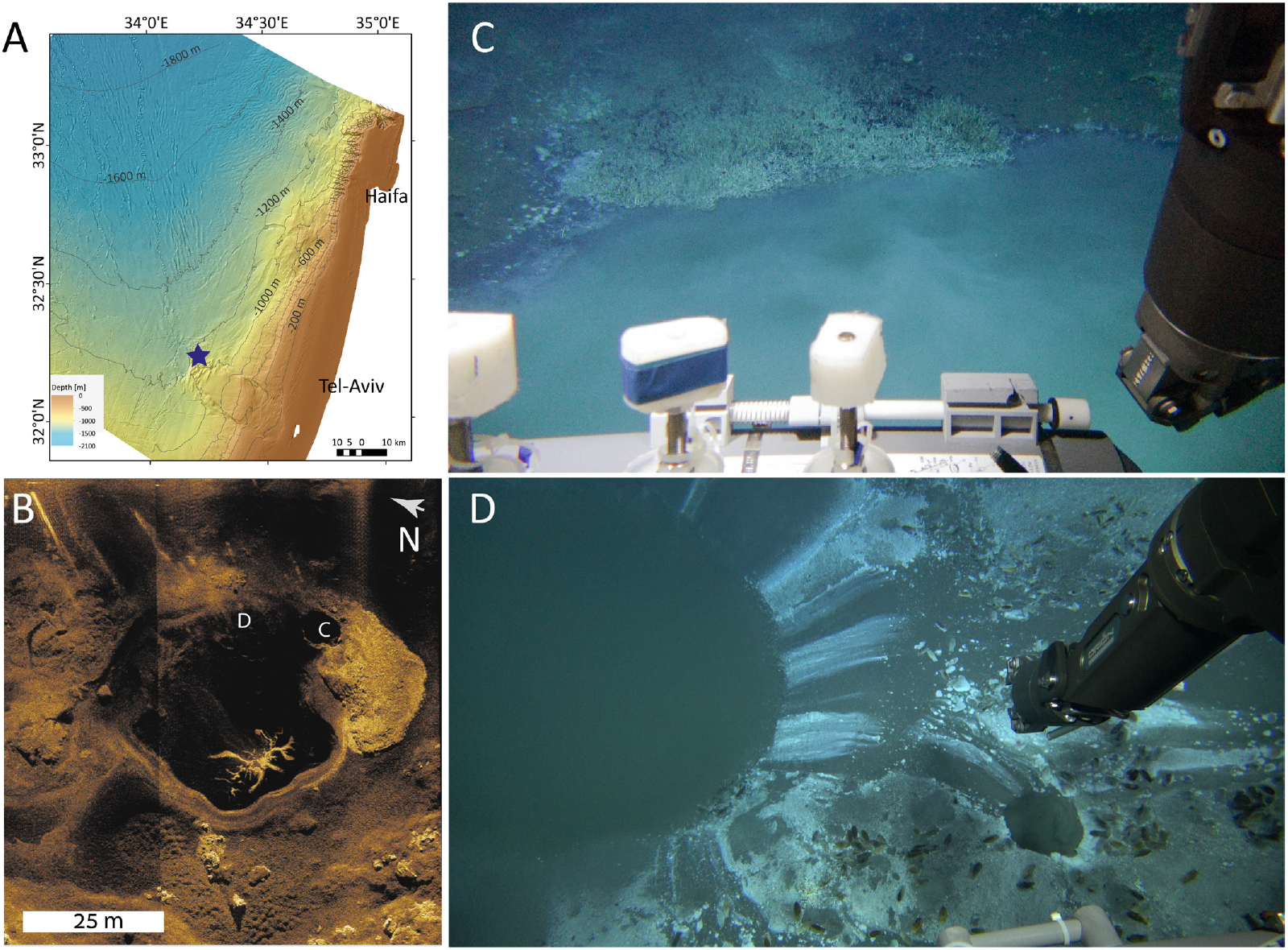
Brine pools at the toe of Palmahim Disturbance, the Southeastern Mediterranean Sea. Bathymetry (A) and synthetic aperture sonar image of the seafloor (B). We sampled primarily the small eastern pool, which is circa 5 m in diameter, and filled with brine (image in panel C, samples BP1-2, 4-5). Gas bubble flares can be seen in the water column. The large pool is only partially filled with brine. A cascade of smaller pools is found at the northern part of the large pool (image in panel D).

### 2.2. Physicochemical parameters

Onboard, water samples were immediately collected directly from the Niskin bottles. The brine salinity (expressed as conductivity), pH, and dissolved oxygen concentrations were measured onboard immediately upon sampling using a WTW Multi 3630 IDS sensor. The pH electrode was calibrated against the standard NBS buffers. Dissolved inorganic nutrients (NOx, NH_4_, PO_4_, Si(OH)_4_) were determined using a Seal auto-analyzer system with colorimetric detection (Sisma-Ventura and Rahav, 2019). The reproducibility of the analyses was determined using certified reference materials (CRM): MOOS 3 (PO_4_, NO_X_, NH_4_), VKI 4.1 (NO_X_), and VKI 4.2 (PO_4_). Results were accepted when measured CRM’s were within ± 5% of the certified values. Total phosphorus (TP) and total nitrogen (TN) concentrations were measured on filtered (Minisart® 0.45 μm) samples following potassium persulfate digestion and ultraviolet (UV) photo-oxidation, using a digestion block system (Seal Analytical). The dissolved organic phosphorous (DOP) and nitrogen (DON) concentrations were determined by subtracting PO_4_/NO_X_ from the total concentrations (TDP and TDN). Quality control of the nutrient measurements over the years was performed with the use of internal and certified reference standards and by participation in international laboratory performance exercises (QUASIMEME). The limit of detection (LOD), estimated as three times the standard deviation of 10 measurements of the blank (low nutrient seawater collected from the off-shore surface SEMS), were 8 nM for PO_4_, 50 nM for total dissolved phosphorus (TDP), 80 nM for NO_2_+NO_3_ (NOx), 90 nM for NH_4_, and 0.74 µmol for total dissolved nitrogen (TDN).

Total Alkalinity (A_T_) was measured by potentiometric titration with a Methrom, 848 Titrino plus system using the Gran method to calculate A_T_ (Ben-Yaakov and Sass, 1977). The A_T_ titration acid was calibrated using certified reference seawater supplied by the Scripps Institution of Oceanography, CA (Dickson et al., 2003). The pH of seawater samples was measured in the total hydrogen scale at a constant temperature of 25.000±0.006 °C with the CONTROS HydroFIA pH flow-through system, based on the ratios of light absorption at 434 and 578 nm before and after adding the factory-calibrated color indicator m-Cresol purple (Aßmann et al., 2011). Dissolved inorganic carbon (DIC) values were calculated from corresponding values of A_T_ and pH using the CO2sys2.1 program (Lewis and Wallace, 1998) and the total hydrogen scale thermodynamic dissociation for the carbonate system in seawater (Millero, 2010). Pore water sulfide concentrations were determined in accordance with the previously-described procedure (Soto et al., 2023)outlined by Soto et al. (2023). Pore water samples were extracted using a Rhizon sampler. Subsequently, an aliquot of each sample was fixed with a 1:10 ratio of a 0.5% (w/vol) zinc acetate solution and immediately frozen until further analysis. Sulfide concentrations were measured using a spectrophotometer, using the methylene blue assay (Cline, 1969).

### 2.3 Microbial cell counts

Brine samples (1.7 ml, BP3-5) were fixed with microscopy-grade glutaraldehyde (1% final concentration, Sigma, G7651), stained with SYBR green (Applied Biosystems cat #S32717) and incubated for 10 min in the dark. Stained samples were enumerated by discrimination based on green fluorescence (530 nm) and side scatter using an Attune® acoustic focusing flow cytometer (Applied Biosystems) (Frank et al., 2017).

### 2.4 Leucine uptake assay

Brine samples BP3 and BP4 were spiked with L-[4,5-^3^H] leucine (Perkin Elmer, specific activity 100 Ci mmol^-1^) at a final concentration of 20 nmol leucine L^-1^ (Kirchman et al., 1985). The samples were incubated in the dark at ∼ 20 °C, fixed using trichloroacetic acid (100%), and treated following the micro-centrifugation protocol (Smith et al., 1992). Disintegrations per minute (DPM) were counted using a TRI-CARB 2100 TR liquid scintillation counter (Packard). Blank background activity in TCA-killed samples was measured. Production rates were determined using a conversion factor of 1.5 kg C mol^-1^ per every mol of leucine incorporated (Sisma-Ventura and Rahav, 2019).

### 2.5 Primary productivity

Carbon fixation rates were estimated in subsamples of brines BP3 and BP4 using the ^14^C incorporation method (Nielsen, 1952), immediately upon recovery. Briefly, collected water samples (45 ml in 50 ml Falcon tubes, with small aerobic headspace) were spiked with 5 µCi of NaH^14^CO_3_ (Perkin Elmer, specific activity 56 mCi mmol^−1^). The samples were incubated for 24 h at ∼ 20 °C in the dark. For the added activity, 5 ml was taken at the beginning of incubation and placed in GF/F filters, and 50 µl of ethanolamine was added to block the purging of radiolabeled bicarbonate from samples. The incubations were terminated by filtering the spiked seawater through GF/F filters (Whatman, 0.7 µm pore size) at low pressure (∼ 50 mmHg). The filters were placed overnight in 5 mL scintillation vials containing 50 µL of 32 % hydrochloric acid to remove excess ^14^C, after which 5 mL of scintillation cocktail (Ultima-Gold) was added. Radioactivity was measured using a TRI-CARB 2100 TR (Packard) liquid scintillation counter.

### 2.6 DNA preservation, extraction, and sequencing

Brines were filtered through either 0.2 µm filters (Supor, Pall) or on GF/F (Pall) until the filter was clogged (tens to hundreds of ml). Filters were kept at −20° C until processed. DNA was extracted from filters with the PowerWater kit (Qiagen, USA), following the manufacturer’s instructions. Metagenomic libraries were constructed using NEBNext® Ultra™ IIDNA Library Prep Kit (Cat No. E7645) and sequenced as circa 100 million 150 bp paired-end reads using the Illumina NovaSeq at Novogene (Singapore).

### 2.7 Bioinformatics

Metagenomic libraries were processed using ATLAS V2.0 (Kieser et al., 2020), with SPAdes V3.14 de-novo assembler (Prjibelski et al., 2020) with --meta -k 21,33,66,99,127 flags. We used metaBAT2 (Kang et al., 2015), VAMB (Nissen et al., 2021) and maxbin2 (Wu et al., 2015) as binners, DAS Tool (Sieber et al., 2018) as a final binner for metagenome-assebled genome (MAG) curation. MAGs were dereplicated with dREP (Olm et al., 2017), using a 0.975 identity cutoff. We used CheckM2 to assess the completeness of MAGs (Chklovski et al., 2023), assigned taxonomy with GTDB-Tk V1.15 (Chaumeil et al., 2020) and annotated them with the SEED and the rapid annotation of microbial genomes using Subsystems Technology (RAST) (Overbeek et al., 2014). Identification of key functions was based on the hidden Markov model (HMM) profiles within METABOLIC (Zhou et al., 2022). *Sulfurimonas* codon treeing was performed based on 500 genes (5 allowed duplications and absences), within the Bacterial and Viral Bioinformatics Resource Center (BV-BRC) framework (Olson et al., 2023), using RaxML version 8.2.11 (Stamatakis, 2014) and MAFFT alignment (Katoh and Standley, 2013). Metagenomic reads were deposited to the NCBI SRA archive under project number PRJNA1042791.

## 3. Results and Discussion

### 3.1 Brines are hotspots of microbial productivity in the oligotrophic SEMS

The brine-water interface of Palmahim brine pools is occupied by dense and productive communities. We determined densities of 0.09±0.03 x 10^6^ cell ml^-1^ in samples BP3-5 (**Figure 2**). These counts are lower than 0.8±0.1 x 10^6^ cell ml^-1^ estimated at the halocline of Lake L’Atalante, which corresponds with the productivity rates of 6.45 µg C L^-1^ d^-1^ (Yakimov et al., 2007). Higher cell counts of 2.1±0.1 x 10^6^ cell ml^-1^ were determined at the upper interface of Lake Thetis (∼3.8 µg C L^-1^ d^-1^ dark productivity rate), reaching 5.6±0.8 x 10^6^ cell ml^-1^at the oxycline, but declining to 0.07±0.01 x 10^6^ cell ml^-1^ in the lower interface (La Cono et al., 2011). In turn, we identified the very high dark productivity rates of 685±16 and 151±24 µg C L^-1^ d^-1^ (BP3 and BP4, respectively), in incubations with air-containing headspace, at temperatures similar to those of in situ ∼20°C (**Figure 2**), resembling the 321.4 µg C L^-1^ d^-1^ rates measured at 24 °C in samples from Crab Spa in the 9°N hydrothermal vent field on the East Pacific Rise (McNichol et al., 2018). These values exceed by two orders of magnitude the background productivity values of circa 0.2 µg C L^-1^ d^-1^ in the deep water of Palmahim Disturbance, those of the productivity in the photic zone (circa 1 µg C L^-1^ d^-1^) and the abovementioned values in L’Atalante and Thethis. The discrepancy between Palmahim and both Thetis and L’atlante rates can potentially represent the differences in in-situ and incubation temperatures (∼20 vs 13 °C, respectively).

**Figure 2:**
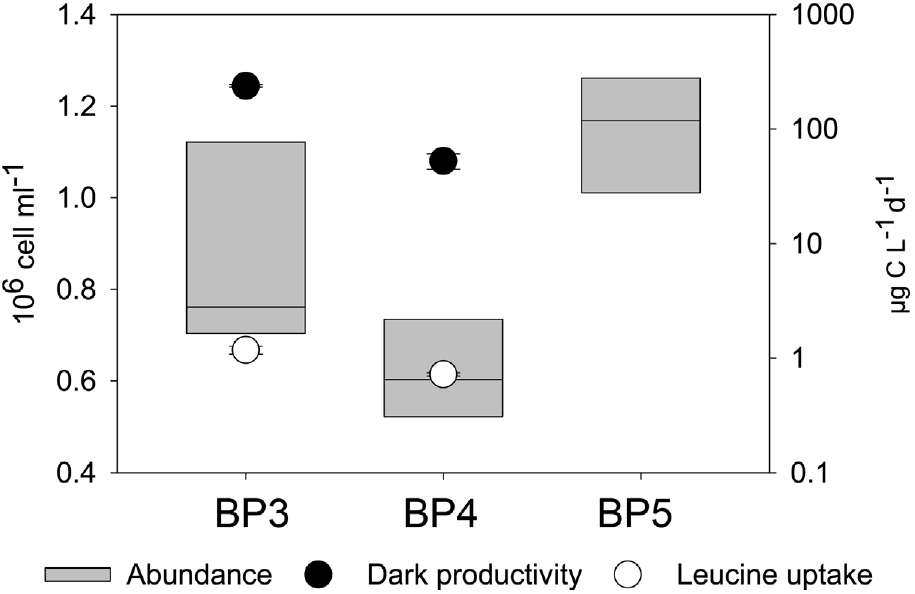
Abundance (left y-axis), primary productivity and leucine uptake rates (right y-axis) of Palmahim brine communities. Background values in the deep water of Palmahim Disturbance are: abundance of ∼ 0.01×10^6^ cell ml^-1^, primary productivity of ∼ 0.2 µg C L^-1^ d^-1^ and leucine uptake rate of ∼0.03 µg C L^-1^ d^-1^.

In turn, the heterotrophic activity measured as leucine uptake was 1.2±0.01 and 0.72±0.02 µg C L^-1^ d^-1^ (BP3 and BP4, respectively), exceeding that of the near-surface waters (∼0.6 µg C L^-1^ d^-1^), and by far that of the 0.03 µg C L^-1^ d^-1^ in the deep sea below 800 m, indicating that secondary productivity is substantial at the brine interface (**Figure 2**). These secondary productivity rates are substantially lower than those of primary ones, hinting that the secondary may be limited by the availability of electron acceptors, as the heterotrophs compete with autotrophs for this resource. Indeed, the lowest maximal growth rates were associated with the autotrophic Campylobacterotra (2±1 h, as opposed to 8±6 h for the whole dataset **Figure 3**).

**Figure 3:**
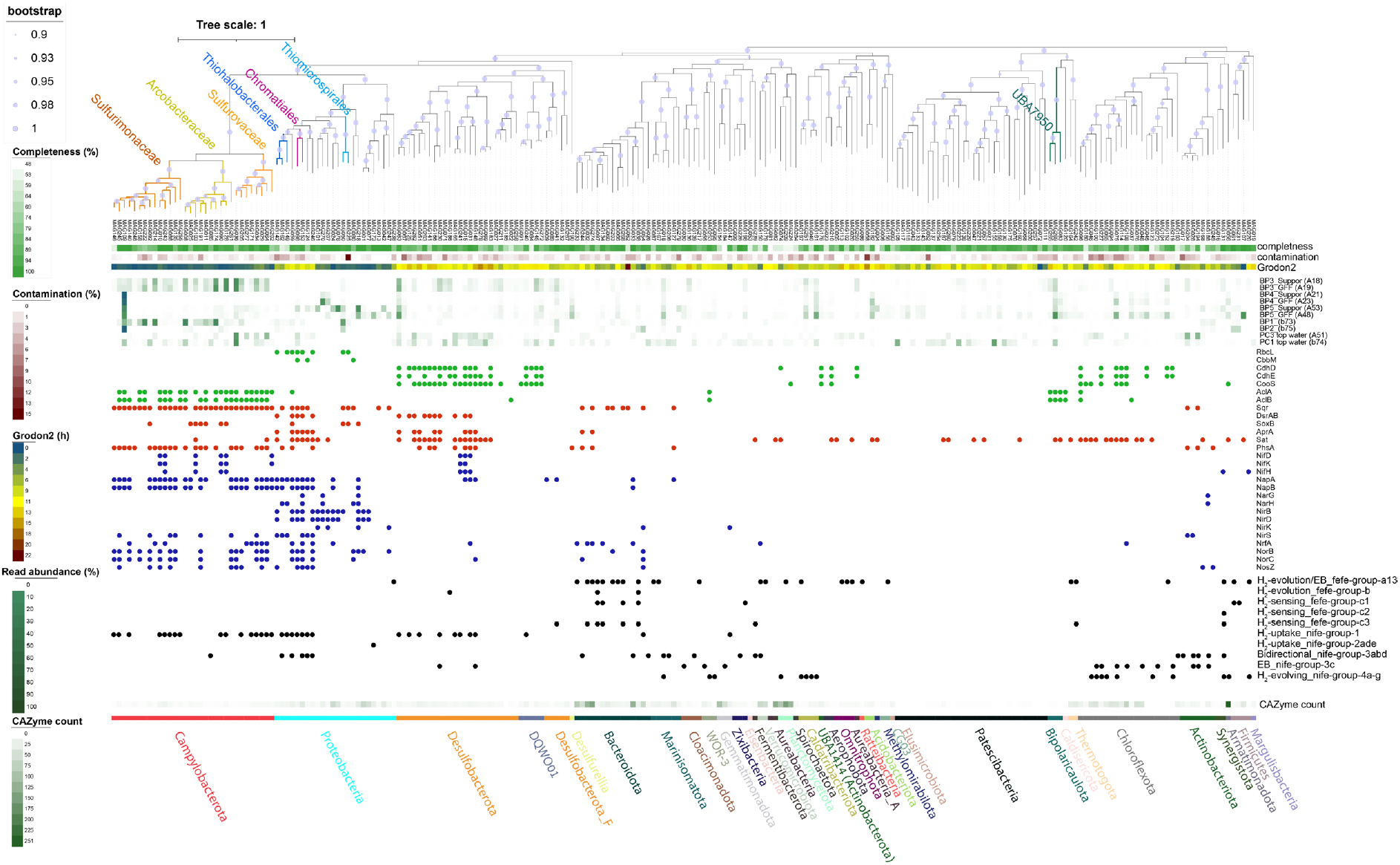
Bacterial metagenome-assembled genomes (MAGs) in Plamahim brines. The GTDB-tk tree is based on the alignment of GTDB-tk bacterial marker gene set. Genome quality stats are based on CheckM2. Minimum replication rates are shown (Grodon2). Key lineages discussed in the text are highlighted. Selected functions in the following modules are presented: carbon fixation (green), dissimilatory S metabolism (red), N metabolism (blue), H metabolism (black). CAZymes are carbohydrate-active enzymes. Electron bifurcation is EB.

The productivity values we measured at the brine interface are in the same order of magnitude as those integrated for the 1150 m water column above, approximated as 365-408 µg C m L^-1^ d^-1^ in the 1150 m deep water column at the Palmahim site (Sisma-Ventura et al., 2022). This highlights the key role of cold seep chemosynthetic productivity in impoverished SEMS. Yet, our estimates with oxygenated headspace may rather represent the potential maximum rates of carbon fixation (see **Supplementary Note 1**).

### 3.2 Specialization at the brine interface limits microbial diversity

Using metagenomics, we curated 266 MAGs (CheckM2 median completeness 92.2%, median contamination 1.4%), spanning 9 archaeal and 39 bacterial phyla (**Supplementary Table ST1)**. The alpha diversity parameters varied among the samples, as some were dominated by a limited number of lineages (Shannon’s H’, based on raw reads that mapped to MAGs was 3±1, **Supplementary Figure S2**). Bacteria and archaea were most diverse in BP1 and BP3, as well as in the top water of the push cores (**Figures 3 and 4**). The microbial communities in samples BP2 and BP4, collected from the same 5 m-wide pool, were less diverse than others due to the dominance of several lineages, such as *Sulfurimonas* (**Figure 3, Supplementary Figure S2**). Given that our sampling likely disturbed the brine interface, it is feasible that we overestimated the diversity in brines, and that the level of specialization in these communities is higher than observed. However, samples BP2 and BP4, and to some extent BP5, likely represent the most pristine brines.

**Figure 4:**
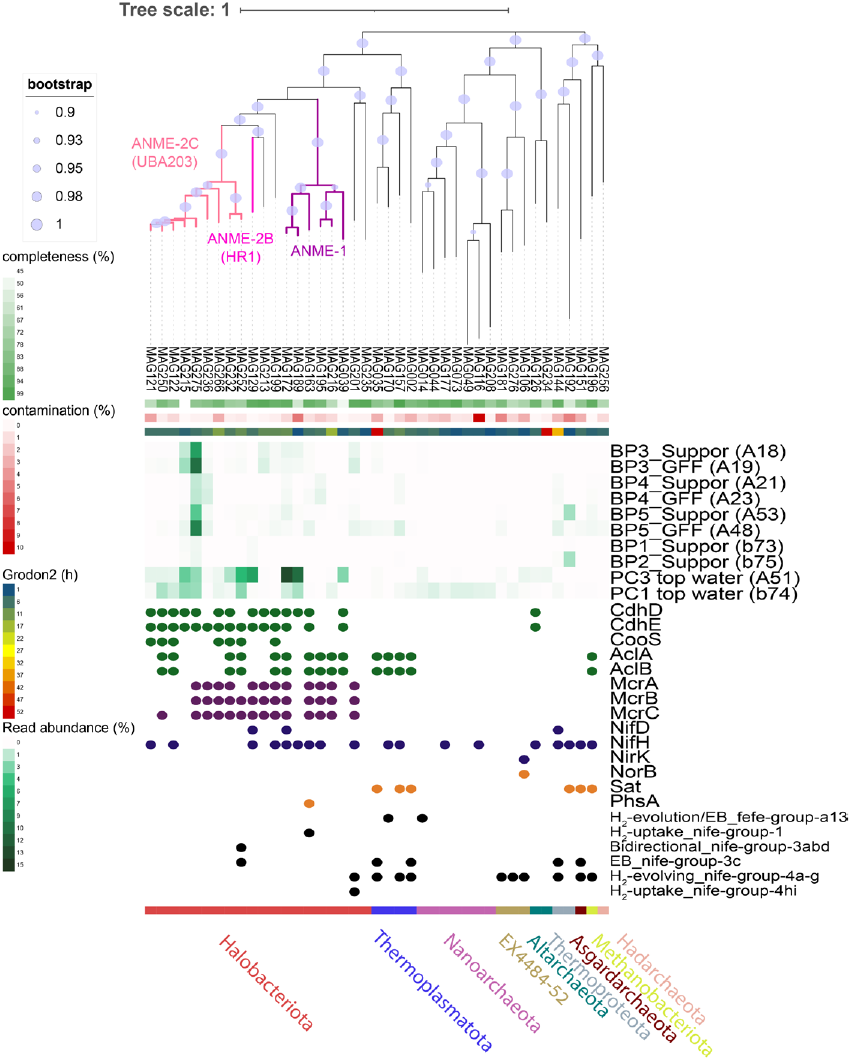
Archaeal metagenome-assembled genomes (MAGs) in Plamahim brines. The GTDB-tk tree is based on the alignment of GTDB-tk archaeal marker gene set. Genome quality stats are based on CheckM2. Minimum replication rates are shown (Grodon2). The putative anaerobic methane oxidizers (ANME) are highlighted. Selected functions in the following modules are presented: carbon fixation (green), methane oxidation (purple), S metabolism (orange), N metabolism (blue), and H metabolism (black).

The communities in the potentially disturbed samples BP1 and BP3 were more similar to those of the seawater overlaying the brine-infused sediments than samples BP2, 4 and 5 (**Supplementary Figure S3**). In turn, the community structure is largely consistent between those filtered on Supor and GF/F filters (0.2 µM and 0.7µm pore size, respectively), suggesting that key lineages do not belong to the 0.2-0.7 µm fraction, and have larger cell sizes (**Supplementary Figure S3**). There are some exceptions. For example, the ANME-2c MAG175 was prominent in BP3 and BP5 and enriched in the GF/F filters that may be selected for ANME aggregates (**Supplementary Figure S2**). In turn, ANME-2c MAG275 (∼1.5% in both filters), represented the most abundant ANME. ANMEs were most diverse in top core water, where ANME-2c, 2b and 1 were present (**Figure 4**), likely due to mixing of the sediment-surface interface by core degassing. We note that ANME-2c were the key organisms in the 0-1 cm layer of PC1, whereas ANME-1 were abundant in the same layer of PC3 (Rubin-Blum et al., in prep.), and the top core water diversity appears to reflect this trend. In turn, a substantial read abundance of Desulfobacterota was found in PC1 (16%), PC3 (13%), BP3 (14% and 15%) and BP5 (6% and 9%) samples. The ETH-SRB1 JACNLL01 MAG063 was particularly common in brine samples BP3 (3.1% and 3.5%) and BP5 (2.5% and 4.5%), likely being the key partner of ANMEs, and potentially mediating metal-dependent anaerobic oxidation of methane (Mn/Fe-AOM) (Xiao et al., 2023). Thus, AOM may contribute to the elevated alkalinity values reaching 15399 μmol kg^-1^ in BP4 (**Table 1**), given the background seawater values of the 2611±9 μmol kg^-1^ (Sisma-Ventura et al., 2022).

**Table 1:** Physicochemical parameters in Palmahim brines. Temp., temperature in-situ; DO, dissolved oxygen; COND, conductivity, A_t_ – alkalinity, TN – total nitrogen, DON – dissolved organic nitrogen.

|  | DENSITY<br>(GR M<sup>-3</sup>) | SALINITY<br>(PSU) | TEMP.<br>(°C) | DO<br>(MG L<sup>-1</sup>) | COND.<br>(MS CM<sup>-1</sup>) | A<sub>t</sub> | PH | PO<sub>4</sub><br>(μM) | NOX<br>(μM) | NH<sub>4</sub><br>(μM) | TN<br>(μM) | DON<br>(μM) | H<sub>2</sub>S<br>(MM) | SO<sub>4</sub><br>(MM) | CH<sub>4</sub><br>(MM) |
| --- | --- | --- | --- | --- | --- | --- | --- | --- | --- | --- | --- | --- | --- | --- | --- |
| BP1 | 1.027 | 39.75 |  |  |  | 3834 | 7.285 | 0.79 | 6.3 | 31 |  |  | 0.6 |  | >2.5 |
| BP2 | 1.027 | 39.78 |  |  |  | 5311 | 7.283 | 1.07 | 5.05 | 30 |  |  | 1.5 |  | >2.5 |
| BP3 | 1.029 | 42.31 | 21.6 | <0.7 | 62.6 | 4705 | 7.312 | 1.10 | 6.72 | 294 | 306 | 6 |  | 29.3 |  |
| BP4 | 1.036 | 54.99 | 21.1 | <0.8 | 77 | 15399 | 7.035 | 6.29 | 2.28 | 1212 | 1574 | 360 |  | 15.2 |  |
| BP5 | 1.029 | 41.82 |  | 0.03 | 62 | 4480 | 7.256 | 1.38 | 7.32 | 242 | 264 | 15 |  | 29.3 |  |
| PC1 | 1.026 | 38.5 |  |  |  |  | 7.5 | 0.09 | 2.52 | 7 |  |  |  |  |  |
| PC2 |  | 44.56 |  |  |  |  |  | 12.73 | 8.35 | 1016 | 1327 | 302 | ~0.6* |  |  |

### 3.3 Campylobacterota plays a key role in carbon fixation

Following previous studies in mud volcanoes in the Mediterranean Sea and in some deep-sea chemosynthetic habitats elsewhere, Campylobacterota, rather than Gammaproteobacteria, were the dominant autotrophs (Felden et al., 2014; Grünke et al., 2011; Inagaki et al., 2004; Omoregie et al., 2008). Campylobacterota was represented in our samples by 32 MAGs, covering Arcobacteraceae, Sulfurimonadaceae and Sulfurovaceae families (**Figure 3)**. Gammaproteobacterial chemoautotrophs, belonging mainly to Thiohalobacterales, Thiomicrospirales and Chromatiales clades were rare, and likely limited to the utmost part of the brine interface, where oxygen is available and carbon fixation via the Calvin-Benson-Bassham cycle is advantageous (e.g., *Thiomicrorhabdus* MAG050 had ∼4% read abundance in BP1, **Figure 3**).

The microbial communities in samples BP2 and BP4, collected from the same 5 m-wide pool, were less diverse than others due to the dominance of several lineages, such as *Sulfurimonas* MAG109 (1.8 Mbp, 99.8% completeness, 0.1% contamination, 184 contigs), which accounted for 67-83% of read abundance in these samples (**Figure 3, Supplementary Figure S2**). MAG109 represents an uncultured lineage, basally branching to the *S. denitrificans* and *S. gotlandica* clade **(Supplementary Figure S4**). In turn, *Sulfurovum* MAG036 (1.73 Mbp, 78.1% completeness, 1% contamination, 331 contigs) was highly abundant in sample BP3 (9% and 13%), as well as in top core seawater (∼5%). Arcobacteriaceae CAIJNA01 MAGs 008 and 174 (2.05 and 2.07 Mbp, 99.8% and 99.95% completeness, both 1% contamination, 245 and 197 contigs) were most abundant in BP1 (22.5% and 20.5% read abundance) but also occurred in BP3 (∼3% and 4-5% read the abundance of MAGs 008 and 174, respectively). We note that these Arcobacteriaceae MAGs represent distinct genotypes, given the 79% average nucleotide identity (ANI). Another Arcobacteriaceae CAIJNA01 MAG128 (2.12 Mbp, 96.7% completeness, 0.7% contamination, 210 contigs) had 6.8% and 8.4% read abundance in BP3. MAG128 also represented a distinct genotype, with 78% ANI to both MAGs 008 and 174. These results likely reflect the distinct niches along the brine-water interface, as well as the level of disturbance during the sample collection. For example, *Sulfurimonas* appear to excel at carbon fixation using nitrate as the terminal electron acceptor, whereas *Sulfurovum* performs best with oxygen (McNichol et al., 2022). We asked which potential adaptive traits underpin the success of these lineages at the brine and sediment-water interface, and if there are genomic clues for separation to distinct niches.

Similar to other chemosynthetic Campylobacterota (Campbell et al., 2006; Nakagawa et al., 2007; Sievert et al., 2008; Wang et al., 2021), these lineages employ the rTCA cycle to fix carbon, as genes encoding the key complexes that catalyze this cycle were found. These include the ATP citrate synthase, OGOR (2-oxoglutarate: ferredoxin oxidoreductase), fumarate reductase and PFOR (pyruvate:ferredoxin oxidoreductase). Some of these organisms are likely obligate autotrophs, as the oxoglutarate dehydrogenase complex is missing, as well as glycolysis and pentose phosphate pathways often appear to be incomplete (Wood et al., 2004). However, Arcobacteriaceae, which in general possessed larger genomes, may be able to take up some organic molecules. For example, TRAP-type C4-dicarboxylate transport system and cluster 3 ABC transporter for basic aa/glutamine/opines were encoded by MAG008, and TRAP dicarboxylate transporter, unknown substrate 6 and L-amino acid ABC transporter were encoded by MAG128. *Sulfurimonas* and *Sulfurovum* likely depend on external sources of vitamin B12 (auxotrophs), lacking cobalamin synthesis clusters that were found in *Arcobacteriacae*, but encoding outer membrane vitamin B12 receptors BtuB. We note that all the abovementioned organisms depend on cobalamin, encoding the B12-dependent methionine synthases (MetH). Thus, we assume that metabolic handoffs are crucial for sustaining the high productivity rates in the brines.

Oxygen and nitrate may serve as electron acceptors at the brine-water interface. Campylobacterota MAGs encoded the high-affinity cbb3-type cytochrome c oxidase CcoNOQP (EC 1.9.3.1), adapted to low oxygen concentrations, is the sole terminal cytochrome c oxidase in all the abovementioned MAGs, indicating the potential to respire trace or detoxify oxygen (Preisig et al., 1996). Both *Sulfurovum* and *Sulfurimonas*, but only MAG008 among the abovementioned Arcobacteriaceae, have the genomic potential for Nap-catalyzed denitrification, as they also encode the high-affinity periplasmic Nap nitrate reductase (*napAGHBF*(*L*) cluster) (Frey et al., 2014; Vetriani et al., 2014). This may reflect the adaptation to low in-situ NOx concentrations of 5.5±2.3 µmol L^-1^ (Wang et al., 1999). Among the abovementioned MAGs, the nitric (NorBC) and nitrous oxide (NosZ) reductases were found only in the *Sulfurovum* MAG036, however, the genetic potential for complete denitrification was widespread in the low-abundance Campylobacterota MAGs (**Figure 3**). One denitrification step, from nitrite to nitric oxide is still obscure, as we found only the ferredoxin-nitrite reductase (*nirA*, EC 1.7.7.1) in *Sulfurovum* and *Sulfurimonas* MAGs.

Which electron donors can be used for chemosynthesis in brine pools? Dihydrogen (H_2_) fuels Campylobacterota in hydrothermal vents (Molari et al., 2023), and could be enriched at the pycnocline, e.g., reaching circa 6 µmol L^-1^ in the Gulf of Mexico brine pool NR1 (Joye et al., 2009). Although considered highly elevated, these H_2_ levels are lower than 162 and 327µM concentrations measured in hydrothermal plumes where the hydrogen oxidizer *S. pluma* was prominent (Molari et al., 2023; Perner et al., 2013). While we did not measure H^2^ directly, metagenomics hints that H_2_ could be locally produced, as we identified diverse, yet often scarce fermenters that can evolve H_2_ via Fe-Fe hydrogenases and belong mainly to Bacteroidota, Marinisomatota, Omnitrophota, Thermotogota and Verrucomicrobiota phyla (**Figure 3**). Thus, an unknown flux of H_2_ may originate from macromolecule fermentation. In turn, the uptake NiFe hydrogenases were widespread among the less abundant Campylobacterota MAGs, as well as in the Arcobacteriaceae CAIJNA01 MAG128 (**Figure 3**). We identified the *hypC,D,E,A* and *F* genes, encoding the hydrogenase metallocenter assembly in *Sulfurimonas* MAG109, so its metabolic potential to use hydrogen remains questionable. In summary, H_2_ is a potential electron donor for primary producers, yet its contribution remains to be determined. Sulfur redox dynamics likely play a key role in the brine interface productivity. Metagenomics suggests that sulfide and thiosulfate can fuel Campylobacterota chemosynthesis, but we should not neglect the disproportionation of inorganic sulfur, which may play a key role in the energy metabolism of these organisms (Slobodkin and Slobodkina, 2019; Wang et al., 2023; Yamamoto and Takai, 2011). As in other Campylobacterota (Wang et al., 2023), the *aprAB*, *dsrAB*, *dsrC*, *dsrMKJOP*, *sat* and *qmoABC* genes of the Dsr and rDSR pathways were not found. Partial or complete Sox systems (clustered *soxXYZAB* genes, as well as a standalone *soxC*) were often present, e.g., in *Sulfurovum* MAG036 and Arcobacteriaceae MAG128, indicating the potential to oxidize thiosulfate (**Table 2**). However, we did not find the Sox system in *Sulfurimonas* MAG109, which appears to have a limited repertoire of sulfur-cycling genes.

**Table 2:** The presence of sulfide: qionone oxidoreductase (Sqr) types, sulfur oxidation (Sox) and metal efflux (FieF) in Palmahim brine Campylobacterota.

| MAG | Family | Genus | Sqr types | Sox | FieF |
| --- | --- | --- | --- | --- | --- |
| MAG261 | Arcobacteraceae |  | II | YZC |  |
| MAG008 | Arcobacteraceae | CAIJNA01 | II | YC | present |
| MAG011 | Arcobacteraceae | CAIJNA01 | II | ABXYZC | present |
| MAG025 | Arcobacteraceae | CAIJNA01 |  | ABXYZC | present |
| MAG085 | Arcobacteraceae | CAIJNA01 | IV | YC | present |
| MAG090 | Arcobacteraceae | CAIJNA01 | II, IV | BXYZC | present |
| MAG103 | Arcobacteraceae | CAIJNA01 | IV | ABYZC |  |
| MAG128 | Arcobacteraceae | CAIJNA01 | II, IV | ABXYZC | present |
| MAG174 | Arcobacteraceae | CAIJNA01 | II | YC | present |
| MAG191 | Arcobacteraceae | CAIJNA01 | II | ABYZC | present |
| MAG255 | Sulfurimonadaceae | GLR69 | II, IV, VI |  | present |
| MAG010 | Sulfurimonadaceae | QNZR01 | II, IV, VI | YZ |  |
| MAG135 | Sulfurimonadaceae | QNZR01 | II, IV |  | present |
| MAG243 | Sulfurimonadaceae | QNZR01 | II, IV, VI |  | present |
| MAG092 | Sulfurimonadaceae | *Sulfurimonas* | II, VI | ABXYZC | present |
| MAG109 | Sulfurimonadaceae | *Sulfurimonas* | II |  | present |
| MAG140 | Sulfurimonadaceae | *Sulfurimonas* | II, IV, VI | C | present |
| MAG146 | Sulfurimonadaceae | *Sulfurimonas* | II, VI | C | present |
| MAG158 | Sulfurimonadaceae | *Sulfurimonas* | II, IV, VI | YZC | present |
| MAG214 | Sulfurimonadaceae | *Sulfurimonas* | II |  |  |
| MAG222 | Sulfurimonadaceae | *Sulfurimonas* | II, IV | BZC | present |
| MAG245 | Sulfurimonadaceae | *Sulfurimonas* | II, IV, VI | YZC | present |
| MAG248 | Sulfurimonadaceae | *Sulfurimonas* | II, IV, VI | YZC |  |
| MAG088 | Sulfurimonadaceae | SZUA-415 | II, VI | YC | present |
| MAG202 | Sulfurovaceae | FS08-3 | II, IV, VI |  |  |
| MAG036 | Sulfurovaceae | *Sulfurovum* | II, IV, VI | ABXYZC |  |
| MAG171 | Sulfurovaceae | *Sulfurovum* | II, IV, VI | C | present |
| MAG205 | Sulfurovaceae | *Sulfurovum* | II, IV, VI | YC |  |
| MAG277 | Sulfurovaceae | *Sulfurovum* | II, IV, VI | ABXYZC |  |
| MAG029 | Sulfurovaceae | SZUA-451 | II, IV, VI | C | present |
| MAG034 | Sulfurovaceae | SZUA-451 | II, IV, VI | C | present |
| MAG064 | Sulfurovaceae | SZUA-451 | II, IV, VI |  | present |

In most Campylobacterota MAGs from Palmahim brines, we identified the canonical sulfide:quinone oxidoreductase (Sqr), needed to oxidize sulfide (Molari et al., 2023), whereas the MAGs differed in Sqr types (**Table 2**). For example, *Sulfurovum* MAG036 encoded type II, IV and VI Sqrs, needed for energy conservation based on oxidation of sulfide (Han and Perner, 2016; Marcia et al., 2010). In turn, *Sulfurimonas* MAG109 encoded only the type II Sqr. Type II Sqr may have a low affinity to sulfide (Km in the millimolar range), whereas type VI is likely adapted to lower concentrations (Han and Perner, 2016). Thus, Sqr diversity may suggest either the ability to cross sulfide gradients (e.g., all the Sulfurovaceae MAGs, encoding type II, IV, and VI Sqrs), or specific adaptation to high sulfide concentrations (e.g., *Sulfurimonas* MAG109). We note that type II Sqr in Campylobacterota MAGs is often linked to a reoccurring gene cluster, potentially involved in the disproportionation of inorganic sulfur. For example, in MAG 109, two copies of the *psrB* gene (polysulfide reductase), two copies of *phsA* gene (thiosulfate reductase/polysulfide reductase) and *nrfD* (polysulfide reductase) follow the *sqr* gene, in similar to the *phsA*/*psrA*, *ttrB*, and *nrfD* cluster in Desulfobacterota that disproportionate inorganic sulfur or reduce thiosulfate, polysulfide, or tetrathionate with an electron donor such as hydrogen (Bell et al., 2020) (**Figure 5**). Together with the widespread occurrence of *phsA* genes in brine Campylobacterota (**Figure 3**), the abovementioned observation provides evidence for the disproportionation of inorganic sulfur by these organisms.

**Figure 5:**
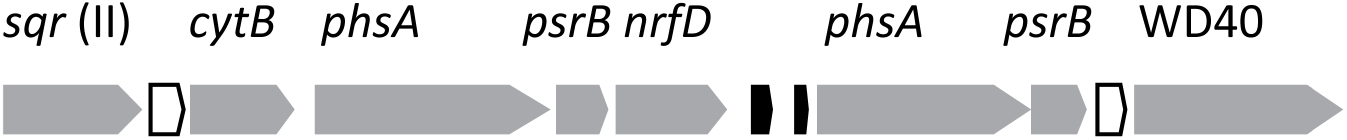
The gene cluster in *Sulfurimonas* MAG109, indicating the genomic potential for inorganic sulfur disproportionation or reduction of thiosulfate, polysulfide, or tetrathionate with dihydrogen as an electron acceptor. Open frames indicate genes that encode putative cytochromes C, black ones encode hypothetical proteins. WD40 is a gene that encodes a WD40 repeat domain-containing protein of unknown function.

We also hypothesized that traits involved in osmoprotection and metal detoxification can be advantageous for Campylobacterota, which inhabit saline brines (∼60 PSU) with elevated concentrations of potentially toxic trace cations, e.g., 3-5 and up to 58.7 µmol L^-1^ Ba^2+^ and Li^+^, respectively (Herut et al., 2022). The use of organic osmoprotectants appears to be limited in brine Campylobacterota, as no MAGs encoded trehalose synthesis, only Sulfurovaceae MAG202 encoded ectoine synthase, Arcobacteriaceae MAGs 085, 011 and 174 encoded the *proVWX* genes encoding the glycine betaine/L-proline transporter and *Sulfurimonas* MAGs 109, 140, 034 and 171 encoded the proline/sodium symporter PutP (the two latter mechanisms may not be involved in osmoregulation) (Christgen and Becker, 2019; Kumar et al., 2020; Wood, 1988). The acute response to high salinities in bacteria is often achieved by the increase in the cytoplasmic K^+^ concentration along with the organic counter ion glutamate (Kindzierski et al., 2017). This process is mediated by various types of transport systems, such as Trk, Ktr and Kef (Kindzierski et al., 2017; Stautz et al., 2021). These systems, especially TrkA and KefA transporters, were widespread among brine Campylobacterota. We identified two adjacent genes encoding potassium TrkA-like transporters in *Sulfurimonas* MAG109. We speculate that adjacent genes, encoding i) glycerol-3-phosphate dehydrogenase [NAD(P)^+^] (EC 1.1.1.94), producing a compatible solute glycerol; ii) CitT sodium-dependent anion transporter and iii) a Dabb family protein, involved in osmotolerance of plants (Gu et al., 2004)), can also be involved in osmoprotection. Metal detoxification mechanisms are less clear, however, FieF divalent cation efflux pumps (Munkelt et al., 2004) were widespread among the brine Campylobaterota (**Table 2**). Thus, the brine Campylobacterota has the metabolic potential for osmoregulation and metal detoxification, yet these traits are not limited to the most widespread lineages.

Why does *Sulfurimonas* MAG109 population prevail over its Campylobacterota counterparts in the pristine, anoxic brines? We hypothesize that its key advantage is genome streamlining to fit the particular ecological niche, given the abovementioned evidence for minimalistic carbon, sulfur and nitrogen pathways. Similar genome streamlining and specialization hallmarks *S. pluma*, a dominant lineage in hydrothermal plumes (Molari et al., 2023). In turn, other Campylobacterota can have a broader habitat range, for example, by encoding several Sqr forms to make use of distinct sulfide concentrations. The presence of nitrogenase machinery in several Campylobacterota (Arcobacteraceae MAGs 090, 103 and 128, *Sulfurimonas* MAGs 010 and 143, as well as Sulfurovaceae MAG202) can pose another burden on the fitness of these populations, given bioavailable nitrogen, e.g. ammonia, is replete in the brines. We note that motility can also play a role in niche selection, as most Sulfurimonadaceae and Arcobacteraceae encoded the flagellum assembly genes, whereas *Sulfurovum* MAG036 did not, similar to some cultivars in this genus (Mino et al., 2014; Xie et al., 2019).

### 3.4 Low-abundance bacteria and archaea in Palmahim brines have versatile metabolic roles

A large variety of low-abundance organisms co-occur with Campylobacterota in Palmahim brines. Some key traits were common in specific phyla. For example, the Woods-Ljungdahl (WL) pathway and Dsr machinery for sulfate reduction were prominent in Desulfobacterota, and some Desulfobulbales lineages were diazotrophs (**Figure 3**). Bacteroidetes often encoded multiple carbohydrate-active enzymes and Fe-Fe hydrogenases, highlighting the potential to ferment complex glycans to dihydrogen (Grondin et al., 2017). As described above, ANMEs were consistently found and encoded the key genes for methane oxidation, and in some cases, diazotrophy (ANME-2c MAGs 232 and 236, ANME-2b HR1 MAG129, **Figure 4**). ANME-2c and 2b appear to be the key producers of vitamin B12 in brines (based on the presence of *cobS* gene, encoding cobalamin synthase), supporting the growth of auxotrophs, such as most Campylobacterota.

We identified some typical extremophile lineages that are often enriched in brines, including *Halomonas* (MAG270, circa ∼8% in the BP4 GF/F sample); Caldatribacteriota JS1 (MAG161, 1-4% in BP4 and BP5, also known as Atribacteria); Bipolaricaulia UBA7950 (MAG269, up to 5% in BP5 sample, also known as OP1 - Acethotermia); Anaerolinae B4-G1 (MAG075, ∼2% on GF/F from BP4), Amicenantes (MAG182, ∼2% on GF/F filters from BP4 and NP5) and Marinisomatota (Glass et al., 2021; Kormas et al., 2015; Liu et al., 2019; Shu and Huang, 2022). *Alcanivorax*, which was recently identified as a key hydrocarbon degrader in hydrothermal plumes (Dede et al., 2023), was also present (1.8% in BP4 GF/F). Most of these organisms appear to have the metabolic potential to degrade macromolecules, either via aerobic pathways, yielding ATP and reducing equivalents via Krebs (TCA) cycle (*Halomonas* and *Alcanivorax*), or ferment to acetate and dihydrogen (See key examples in **Figure 6**). The TCA cycle was usually incomplete in anaerobes, which often employed the Woods-Ljungdahl pathway alongside interconversions between pyruvate, acetyl-CoA, formate and acetate (Kremp et al., 2022; Lü et al., 2012; Mühlbauer et al., 2021; Pettinato et al., 2022). These bacteria are likely involved in necromass degradation, as they have the metabolic potential to ferment various sugars, as well as nitrogen-rich compounds such as peptides, amino acids and nucleosides, given the marked abundance of respective transporters (**Figure 6**). They encoded multiple peptidases (particularly abundant in aerobes), as well as CAZymes, potentially to break down necromass macromolecules (**Supplementary Table ST2**), as suggested for other seep and vent systems (Pérez Castro et al., 2021; Zhao et al., 2020). This process may lead to the marked enrichment of ammonium, DON (e.g., the millimolar NH_4+_ levels in BP4 or PC2, DON concentrations above 300 µmol L^-1^, and the high, over 1200, NH_4+_+NO_x_/PO_4-3_ ratio, which is higher by a factor of ∼45 than the bottom seawater, **Table 1**). Alongside subsurface sources, this microbial diagenesis of organic matter is a potential source of inorganic phosphate, reaching high levels of ∼6 µmol L^-1^ in BP4 (**Table 1**), which are still considerably lower than ∼ 40 µmol L^-1^ measured in the Northern Gulf of Mexico brine pools (Bowles et al., 2016), but higher by a factor of 20 than that of the bottom seawater (Sisma-Ventura et al., 2022). These excess nutrients may support the high productivity rates at the brine chemocline.

**Figure 6:**
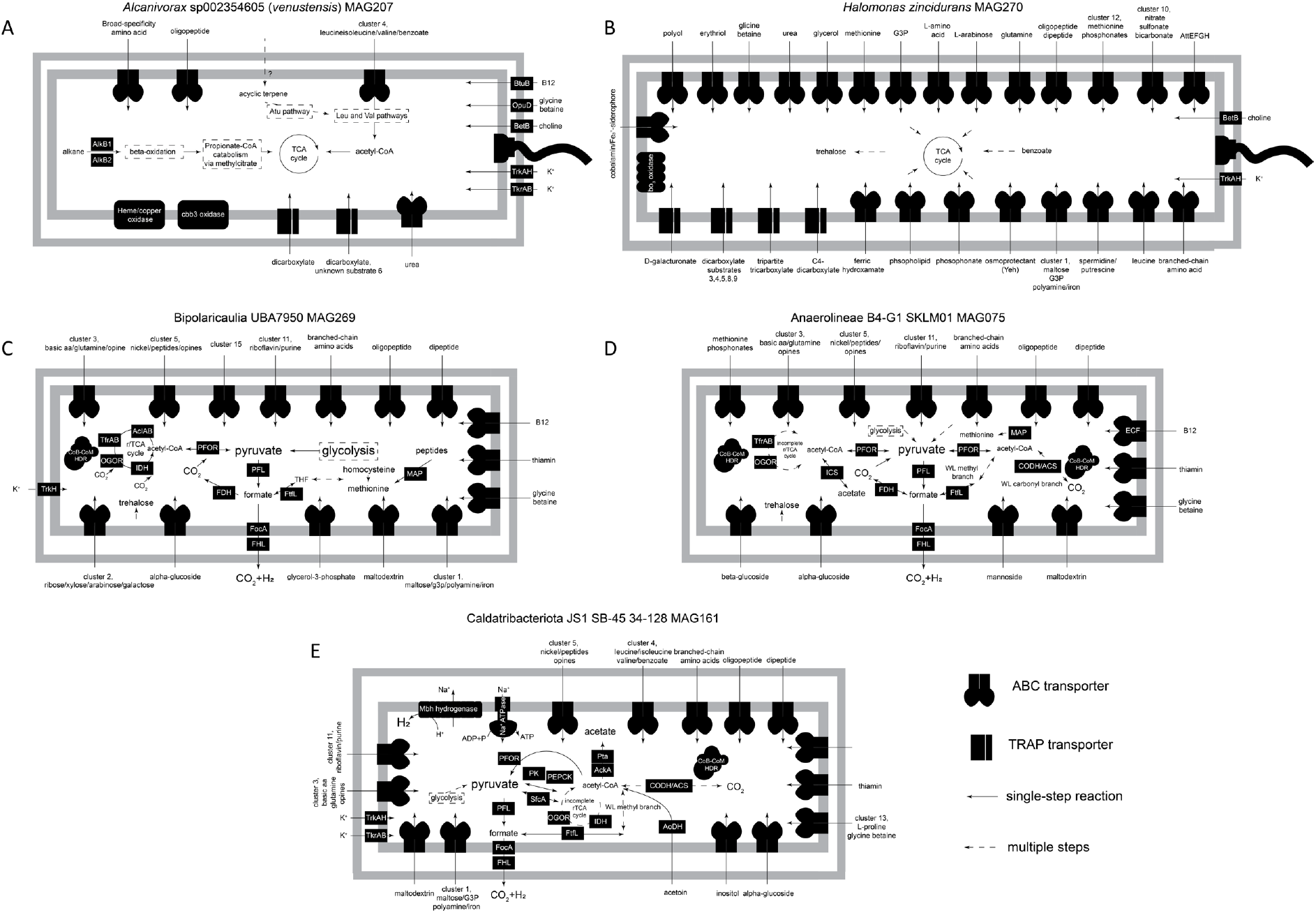
Key metabolic features of aerobic (A and B) and anaerobic (C-E) heterotrophs associated with primary producers in Palmahim brines, based on SEED annotations and manual curation. The following enzymes are abbreviated: OGOR (2-oxoglutarate: ferredoxin oxidoreductase), PFOR (pyruvate: ferredoxin oxidoreductase); IDH (isocitrate dehydrogenase); FDH (formate dehydrogenase); PFL (pyruvate-formate lyase); FHL (formate-hydrogen lyase); ICS (acetyl-CoA synthetase); MAP (methionine aminopeptidase); PK (pyruvate kinase); PEPCK (phosphoenolpyruvate carboxykinase); HDR (heterodisulfide reductase); CODH/ACS (carbon monoxide dehydrogenase/acetyl-CoA synthase). Aerobes import and degrade multiple compounds, which fuel Krebs (TCA) cycle. Anaerobs ferment multiple molecules, and all may produce H_2_, either via formate oxidation or using the Mbh Na-translocating hydrogenase (Caldatribacteriota -JS1, Atribacteria). Caldatribacteriota and Anaeorlineae may produce or consume acetate, and encode the Woods-Ljungdahl pathway. Bipolaricaulia encodes the complete reverse tricarboxylic acid cycle, indicating the potential for ferredoxin-driven carbon fixation. Only aerobes encode genes for the assembly of flagella. The metabolic reconstruction of Amicenantes and Marinisomatota is pending due to poor annotation of proteins in these lineages, however, large numbers of organics transporters were found and putative fermentative metabolism was also estimated in these organisms.

Our observations confirm the genomic potential of Bipolaricaulia UBA7950 (MAG269, as well as the less abundant populations represented by MAGs 083 and 265) to fix carbon via the rTCA cycle, as they encoded all the necessary genes, in particular, type I ATP-citrate lyase (Youssef et al., 2019). The WL pathway, often encountered in Bipolaricaulia (Youssef et al., 2019), was not found. The rTCA cycle in this lineage can be driven by flavin-based electron bifurcation, given the presence of gene cluster encoding OGOR, the Hdr-Flx complex, and thiol-fumarate reductase TfrAB (Rubin-blum et al., 2019). The role and energy budget of the rTCA cycle in UBA7950 metabolism remains to be elucidated.

## 4. Conclusions

Our data suggest that Palmahim Disturbance brines are inhabited by microbial populations comprised of specialists, such as niche-adapted Campylobacterota autotrophs and ANMEs, and diverse, but less dominant generalist heterotrophs. The latter is likely a crucial component of the system, needed for the necromass recycling to produce small organic molecules and H_2_. We only investigated the metabolic potential of several representative heterotrophs, but further work is needed to assess the full complexity of the microbiome at the brine and brine-infused sediment interface. We highlight the importance of metabolic handoffs, for example, the potential dependence of key producers on cobalamin from other organisms, such as ANMEs. However, most syntrophic associations, as well as other biotic interactions, including those with phages, remain to be determined.

## Author contributions

MR-B, YM and BH conceived this study and acquired funding. YM performed geophysical surveys identifying the sampling sites and led the ROV work. GS-V, GA and BH analyzed the chemical parameters. ER and NB estimated microbial abundance and productivity. MR-B performed molecular and bioinformatics analyses. MR-B wrote the paper with the contributions of all co-authors.

## Acknowledgments

The authors thank all individuals who helped during the expeditions, including onboard technical and scientific personnel, and the captains and crew of the E/Vs Bat Galim. We thank Ben Herzberg, Samuel Cohen and Oded Ezra, who operated Yona ROV.

## Data availability statement

The raw metagenomic reads and metagenome-assembled genomes are available on NCBI as BioProject PRJNA1042791.

## Supplementary Information

### Active microbial communities facilitate carbon turnover in brine pools found in the deep Southeastern Mediterranean Sea

Maxim Rubin-Blum, Yizhaq Makovsky, Eyal Rahav, Natalia Belkin, Gilad Antler, Guy Sisma-Ventura, Barak Herut

#### Supplementary Note 1

Given the 0.09±0.03 x 10^6^ cell ml^-1^ counts, the cell-specific activity appears to be as high as 2882±697 and 852±39 fg C cell^-1^ d^-1^, in samples BP3 and BP4, respectively. These values correspond to 912-960 fmol C cell^−1^ d^−1^ rates, measured in the photosynthetic *Aphanizomenon* sp. (Ploug et al., 2010). In our samples, Campylobacterota were the key autotrophs, accounting for approximately 50 and 75% of prokaryotes in these two pools, respectively (**Main Text, Figure 3**). Yet, the rates we measured markedly exceed 23.6±8.5 and 78.4±25.3 fg C cell^-1^ d^-1^ estimated experimentally in cultivated and freshly-collected Campylobacterota from hydrothermal vents (McNichol et al., 2022, 2018). We could underestimate the abundance of autotrophs in the incubations, as microbial communities change during the incubations, and Campylobacterota may become more dominant (McNichol et al., 2022), but productivity in our incubations could have been boosted by the excess oxygen in the headspace. Thus, our estimates may rather represent the maximum potential of carbon fixation.

On the other hand, high productivity rates appear to be necessary to fulfill the estimated carbon need of the quickly-dividing Campylobacterota. For example, Grodon2 interpretation of codon usage patterns indicates the minimal doubling time of 2±1 h for Campylobacterota MAGs curated in this study (**Main Text, Figure 3**). The elongated cells of the *Sulfurimonas* can exceed 5 µm in length, and thus likely incorporate more carbon than the average deep-sea bacterium that is expected to contain roughly 20-34 fg C cell^-1^ (Molari et al., 2013). To sustain the duplication rates of 2h, and considering the high values of 34 fg C cell^-1^, a bacterium may fix 17 fg C cell^-1^ h^-1^ (408 fg C cell^-1^ d^-1^), not considering the loss of secreted carbon. Thus, high cell-specific carbon fixation rates are feasible in Camplylobacterota.

## Supplementary Excel files

**Table ST1:** Taxonomy, completeness and read abundance of metagenome-assembled genomes in this study.

**Table ST2:** METABOLIC reconstruction of metabolism in metagenome-assembled genomes curated in this study.

**Supplementary Video:** Remotely-operated vehicle sampling of Palmahim Disturbance brines and sediment push cores.

---

**Figure S1:**
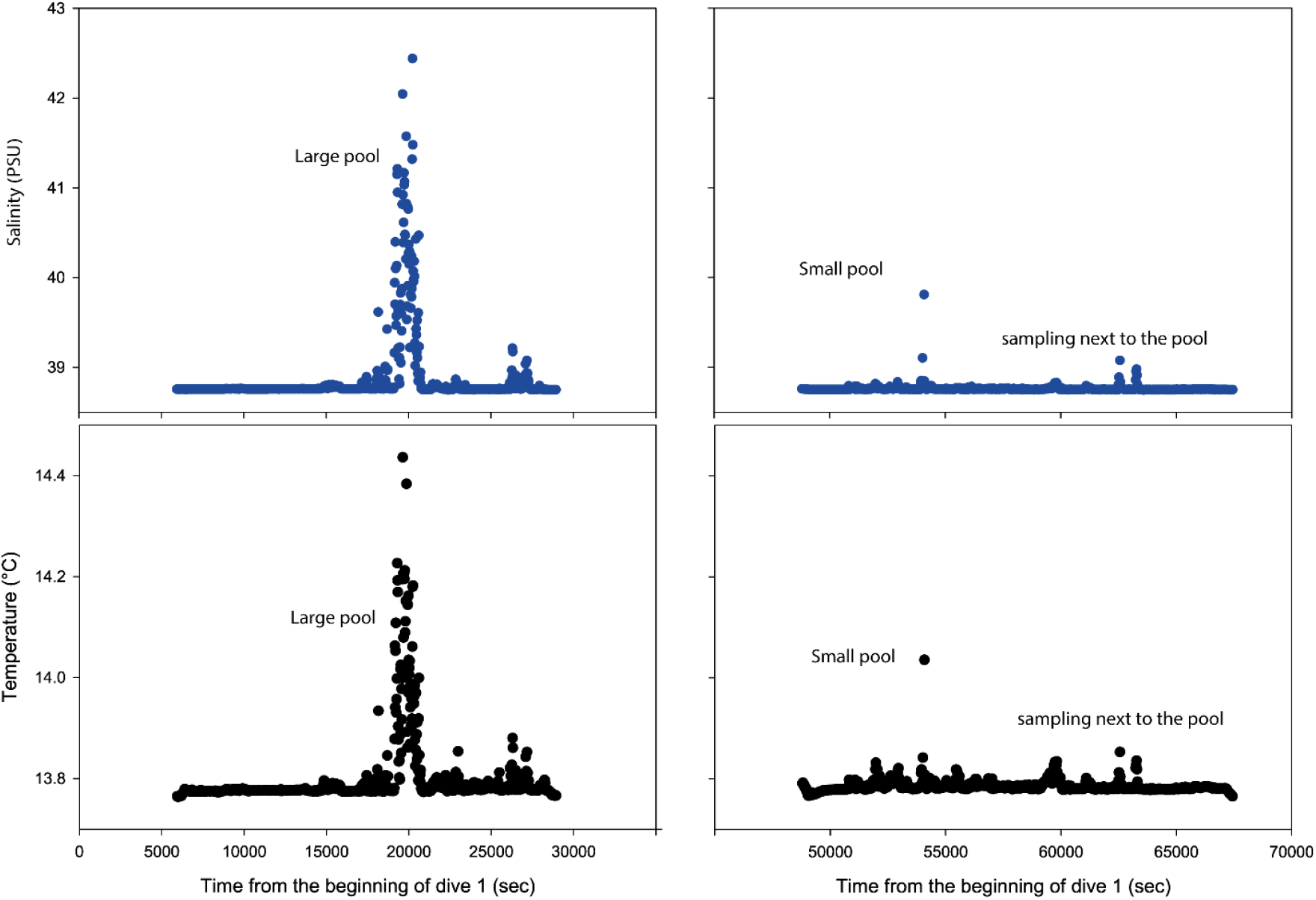
CTD measurements during two ROV dives in April 2021 to Palmahim Disturbance. The CTD was mounted at the level of the Niskin bottle. Temperature and salinity peaks represent the ROV sampling.

**Figure S2:**
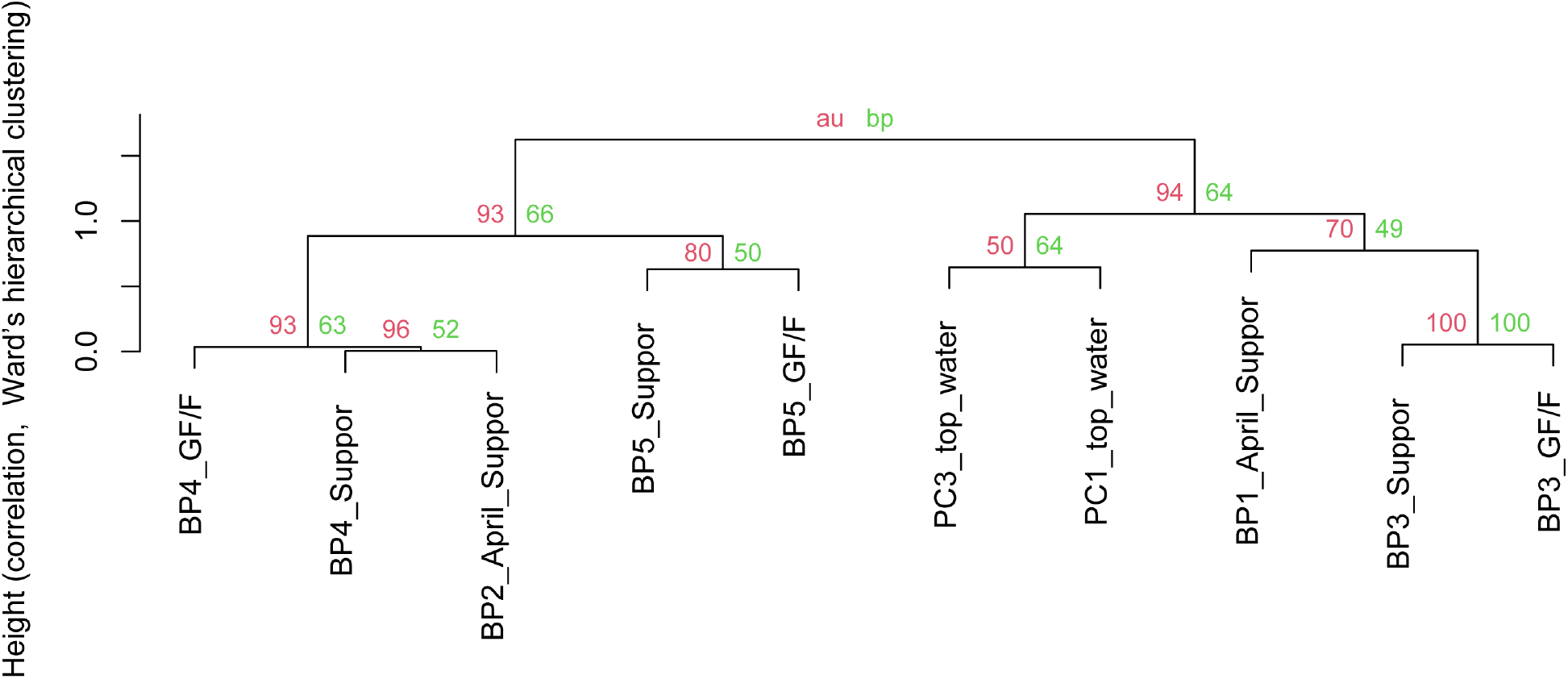
Ward’s hierarchical clustering of samples using pvclust R package, based on metagenomic read abundance. Approximately unbiased p-value (au, red) and bootstrap probability (bp, green) are shown next to branches.

**Figure S3:**
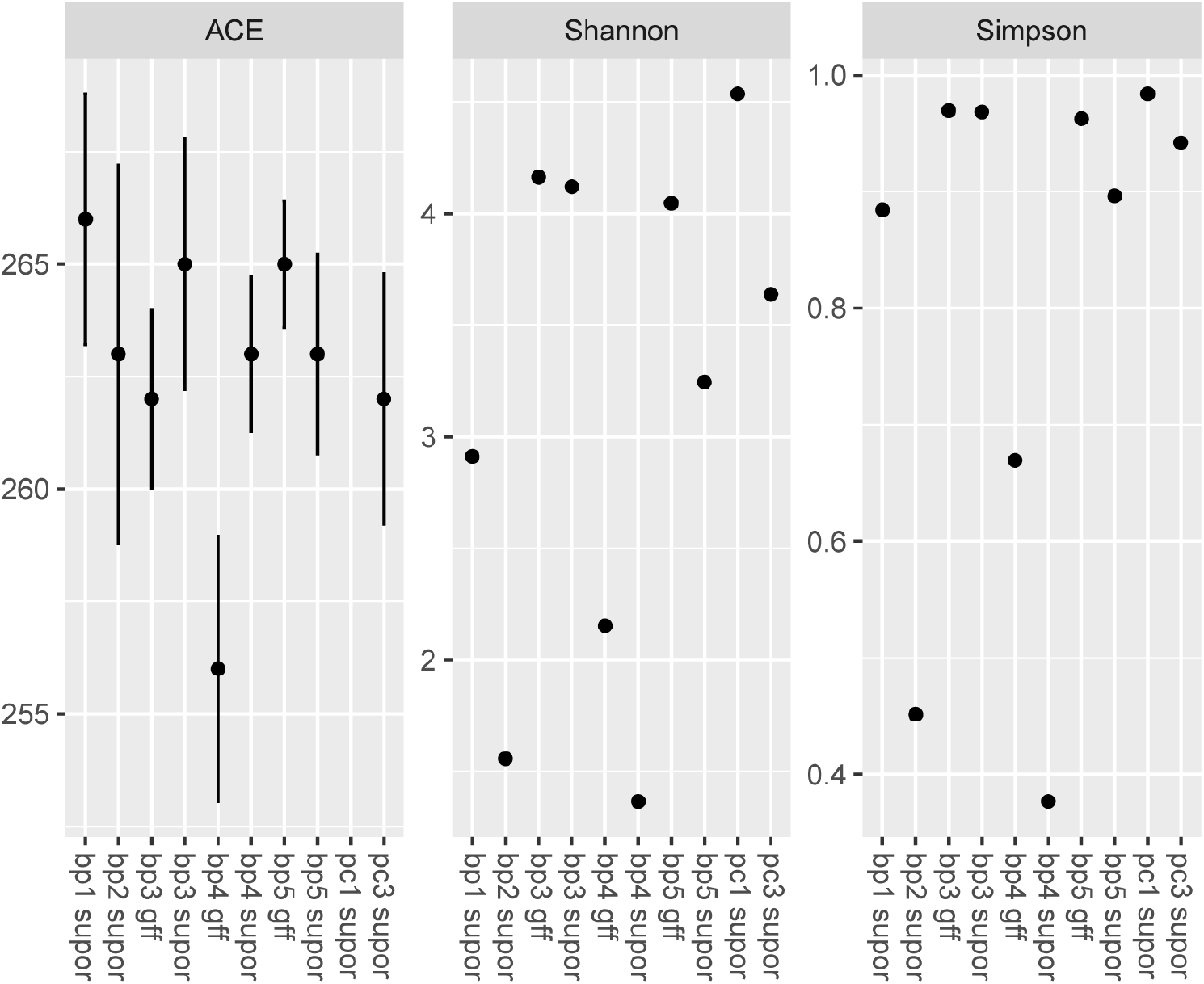
Alpha diversity parameters based on read abundance of metagenome-assembled genomes.

**Figure S4:**
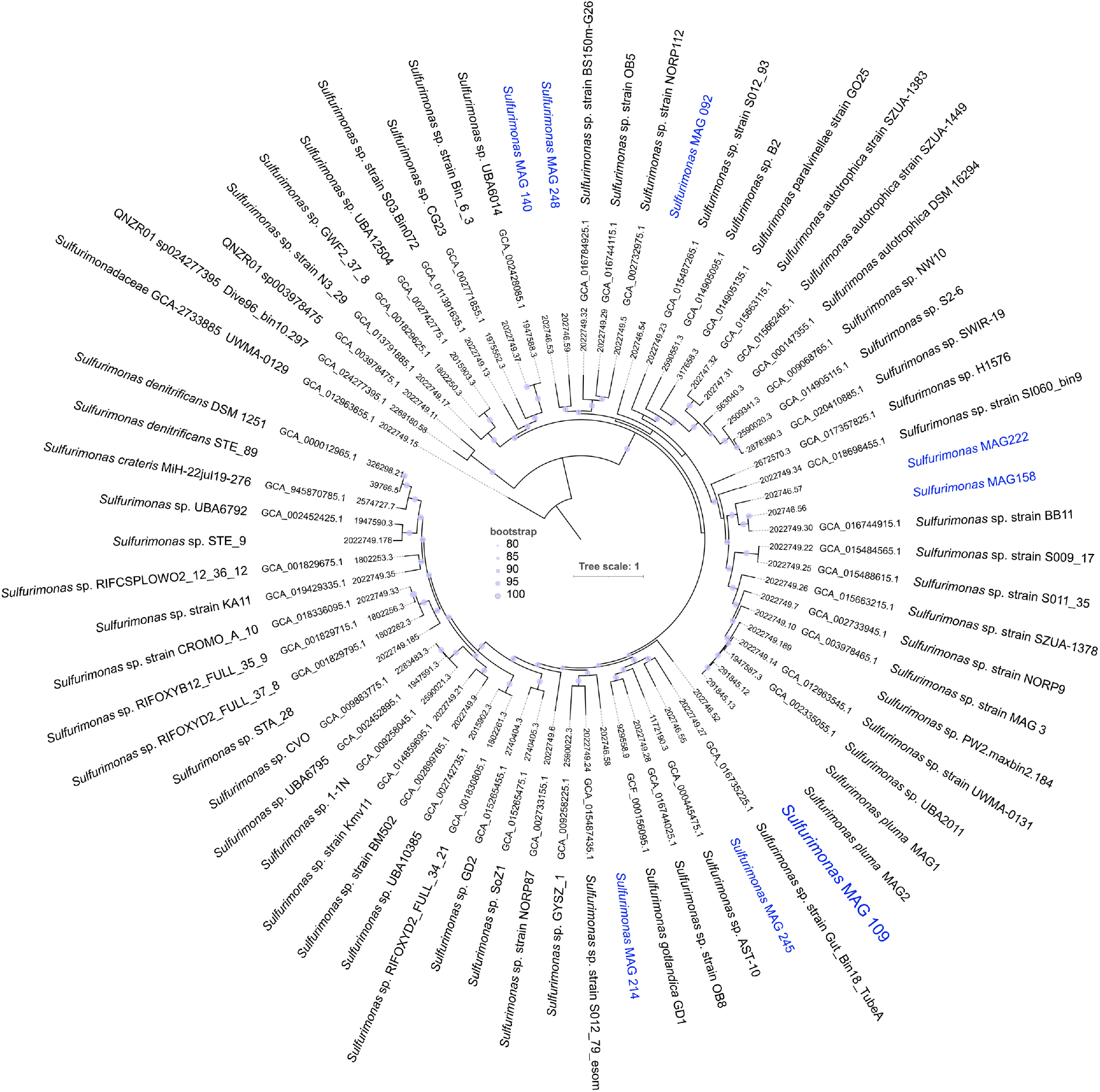
Codon phylogenomic tree of *Sulfurimonas* clade, showing the placement of metagenome-assembled genomes in Palmahim brines. Sulfurimonadaceae QNZR01 genomes are used as the outgroup. BV-BRC and GeneBank accession numbers are shown next to the respective genomes, where available. The tree scale represents the number of substitutions per site. Palmahim (blue) lineages are highlighted.

**Figure S5:**
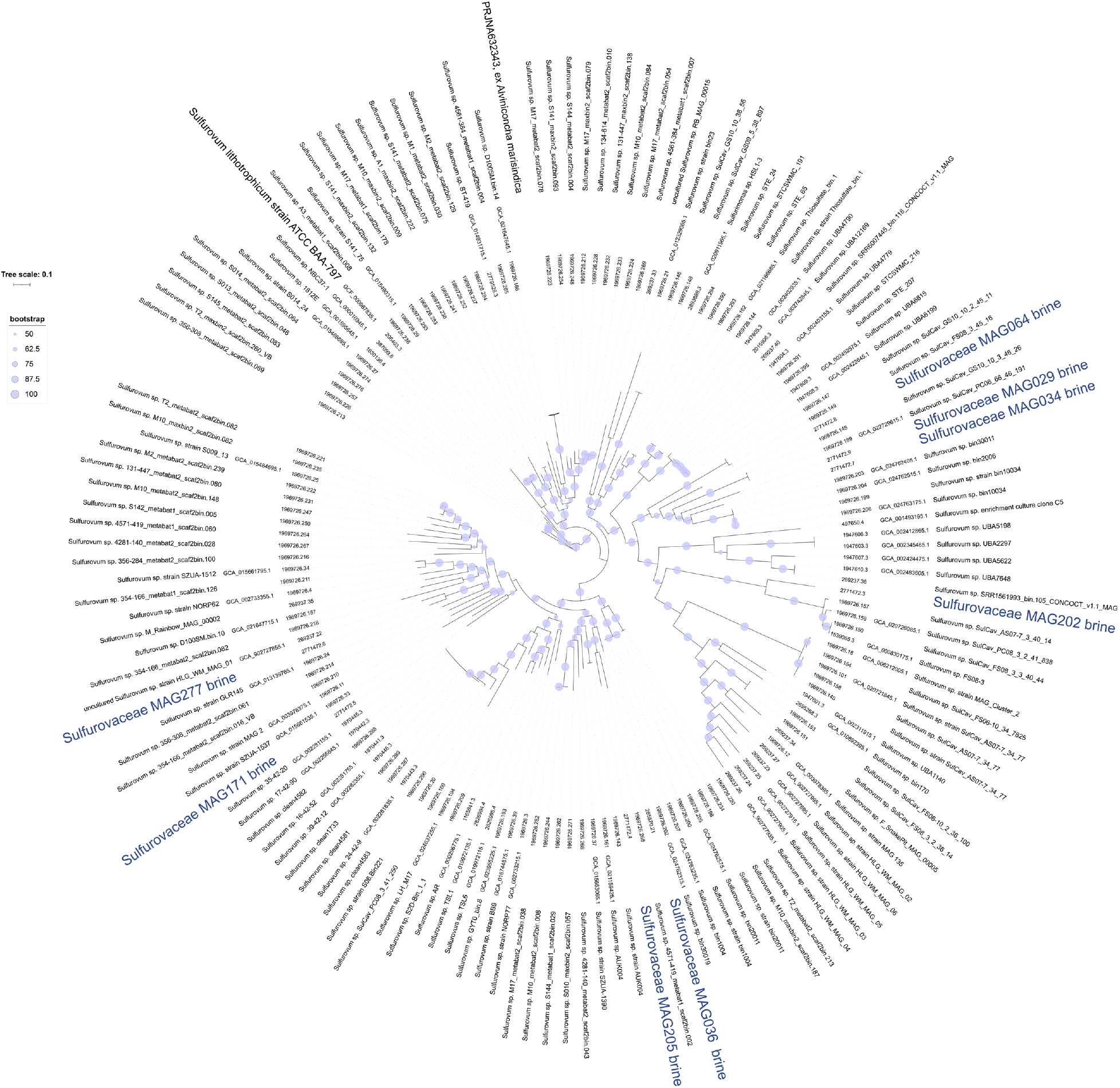
Codon phylogenomic tree of Sulfurovaceae clade, showing the placement of metagenome-assembled genomes in Palmahim brines. The tree is unrooted. BV-BRC and GeneBank accession numbers are shown next to the respective genomes, where available. The tree scale represents the number of substitutions per site. Palmahim (blue) and representative lineages are highlighted.

**Figure S6:**
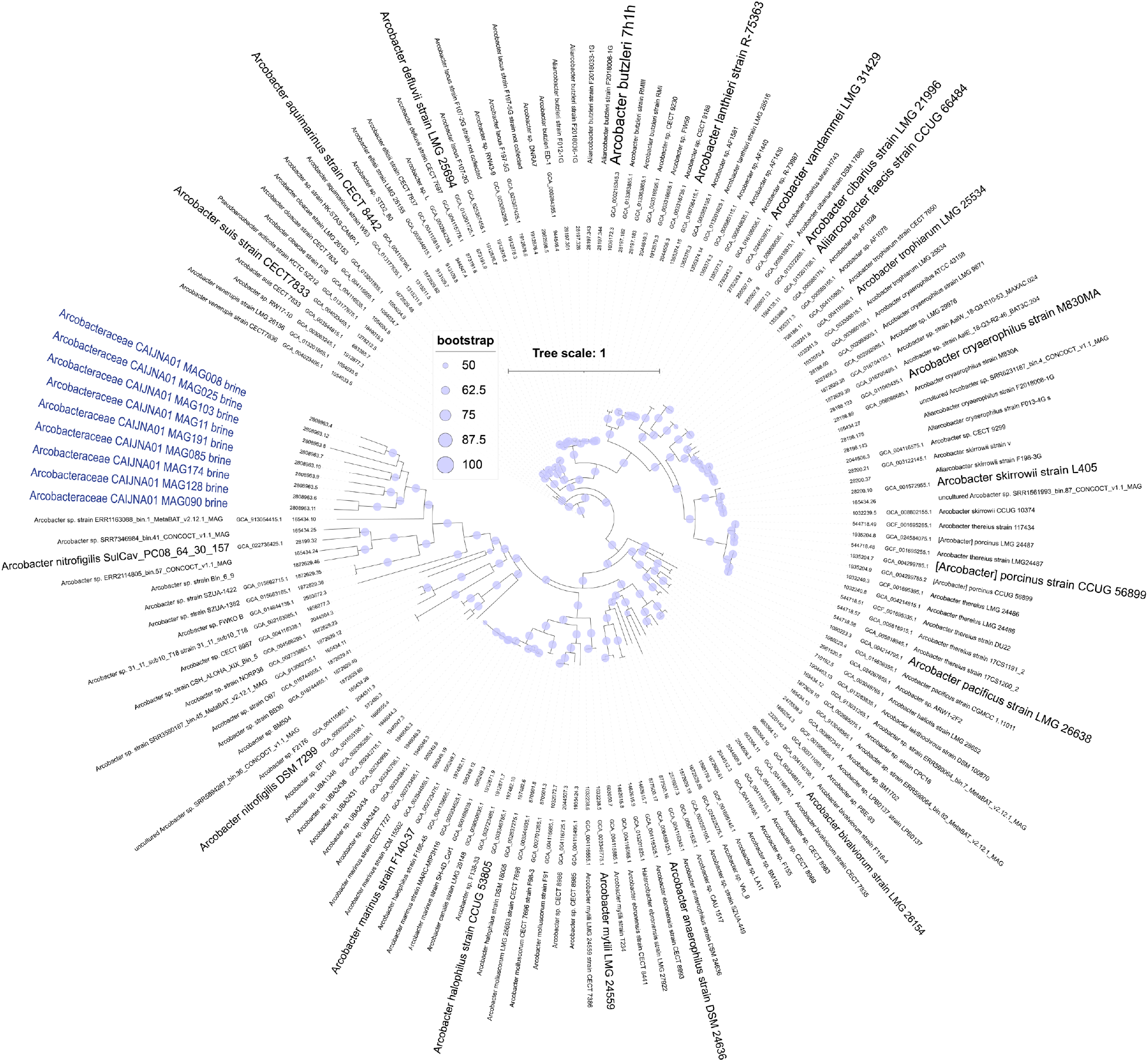
Codon phylogenomic tree of Arcobareriaceae clade, showing the placement of metagenome-assembled genomes in Palmahim brines. The tree is unrooted. BV-BRC and GeneBank accession numbers are shown next to the respective genomes, where available. The tree scale represents the number of substitutions per site. Palmahim (blue) and representative lineages are highlighted.

